# Harlequin Frogs (*Lysapsus*) prefer mild places to park

**DOI:** 10.1101/2020.06.11.146613

**Authors:** Carolina Cunha Ganci, Zaida Ortega, Diogo B. Provete

## Abstract

Temperature affects most aspects of ectotherms’ life history, including physiology and behavior. Studying thermal sensitivity of jumping performance in frogs can help understanding the influence of temperature on different aspects of frog life. Still, studies on the effects of temperature on amphibians are commonly carried out on terrestrial and tree species, creating a gap for aquatic species. We experimentally tested the thermal sensitivity of jumping performance of the Uruguay Harlequin Frog, *Lysapsus limellum*, assessing three measures: response time, distance of first jump, and total distance travelled. We hypothesized that individuals submitted to extreme temperatures would increase response time, decrease first jump distance, and increase total jump distance. We used an arena with a gradient of air temperature (T_a_) ranging from 20 to 40 °C. We placed frogs at different T_a_ and stimulated them to jump. Then, we analysed the influence of T_a_ on the three estimates of jumping performance, using generalized additive models. We found that temperature affected all three measurements of jumping performance, but some relationships were stronger than others. Extreme temperatures increased response time, reduced first jump distance, and increased total distance. The effect was weaker for response time and first jump distance, but substantially stronger for total distance jumped. Although individuals under extreme temperatures experience a reduced jumping performance, they travelled longer distances to find areas with milder temperatures. Thus, we showed that *L. limellum* thermoregulates by means of behavior, moving through places at different thermal conditions. Additionally, benefits of displacing to thermally suitable places -in terms of enhanced jumping performance-are bigger than the costs of jumping at reduced locomotor performance, at least under experimental conditions. Our results can help understand how climate change affects the locomotor performance of Neotropical amphibians.

## 1. Introduction

Temperature is one of the main environmental variables driving the ecology and evolution of organisms (Angilletta, 2009), since it influences the free energy available to drive reactions forward (Little & Seebacher, 2016). Temperature can affect animals in two ways. Firstly, through a top-down effect, when temperature variation is perceived by the central nervous system and modulates behavior. Secondly, through a bottom-up effect, in which temperature variation changes biochemical reaction rates, which alters multiple cellular and systemic processes that consequently affect the organism as a whole (Abram et al., 2017).

Thermal ecology includes two main aspects: thermal sensitivity (i.e. the dependence of physiological performance on temperature) and thermoregulation (i.e. the maintenance of relatively stable body temperatures despite environmental variation). Ectotherms, that is, individuals that depend on environmental heat sources for thermal regulation (Huey & Stevenson, 1979; Angilletta, 2009), show a gradient of performance on thermoregulation, from ideal perfect thermoregulators, whose body temperature are always constant regardless of the environment to thermoconformers whose body temperatures always match environmental temperatures. Additionally, the thermal sensitivity of ectotherms varies from thermal generalists, which perform well under a broad range of environmental temperature, to thermal specialists, whose performance is suitable only in a narrow temperature range (Angilletta, 2009). Consequently, environmental temperature influences most aspects of physiology and behavior of ectotherms, such as foraging ability (e.g., Dell et al., 2014; Halliday & Blouin-Demers, 2016), immune function (e.g., Wright & Cooper, 1981; Mondal & Rai, 2001), rates of feeding and growth (e.g., Huey & Stevenson, 1979; Dastansara et al., 2017) or locomotion (Whitehead et al., 1989; Knowles & Weigl, 1990; Lai et al., 2018). Finally, environmental temperature modulates locomotor performance, which is tightly linked with survival and consequently to fitness of individuals and populations (Huey & Berrigan, 2001; Husak et al., 2006; Seebacher & Walter, 2012).

As ectotherms, anuran amphibians are temperature sensitive organisms. Thus, studying the thermal dependency of their locomotor performance allows us to understand how temperature affects the whole-animal activity, linking ecology and physiology (Huey & Stevenson, 1979). From an ecological perspective, assessing thermal sensitivity of locomotion provides valuable information on how temperature influences prey capture, predator avoidance, or social interactions (Huey & Stevenson, 1979). Different locomotory strategies allow anurans to live in several environments, such as arboreal, fossorial, ground dwelling, or aquatic (e.g., Emerson, 1985; Duellman & Trueb, 1986; Gomes et al., 2009; Jorgensen & Reilly, 2013; Citadini et al., 2018). Therefore, predator scape strategies depend on morphological aspects that were under different selective regimes according to the environment in which each species lives (Emerson, 1978; Zug, 1978). For example, fossorial and terrestrial species have relatively short hind limbs which give them lower jumping performance compared to aquatic species (Zug, 1978; Citadini et al., 2018; Rebelo & Measey, 2019). For aquatic frogs, jumping is the main strategy for escaping predators, since swimming is more costly, because water is 800 times denser and 60 times more viscous than air, making swimming movements harder than jumping (Nauwelaerts et al., 2005). However, most previous studies testing the effect of temperature on jumping performance of frogs used terrestrial and arboreal species (i.e. Hirano & Rome, 1984; Whitehead et al., 1989; Navas et al., 1999). Therefore, little is known about how fully aquatic species perform along a thermal gradient.

Jumping performance depends on environmental temperature in a curvilinear relationship. This is usually assessed by a thermal performance curve (Huey & Stevenson, 1979; Angilletta, 2009). The most commonly used are optimal temperature (the temperature which maximizes performance), thermal performance breadth (the temperature range comprising an arbitrary performance threshold, normally 80%), and thermal limits (the temperature range for which performance is zero) (Angilletta, 2009; Vitt & Caldwell, 2013). These concepts can be connected by the morphology-performance-fitness diagram (Arnold, 1983; 2003). This conceptual construct is useful to link data from experiments measuring performance with morphological measurements to understand its consequences on fitness. According to Navas et al. (2008), temperature can affect muscle performance in several ways and intensities, varying among species, but locomotor performance is maximum under low to moderate temperatures and becomes impaired at higher temperatures. Accordingly, fitness itself is usually understood as a variable measured at the level individuals (Orr, 2009). However, it may also be estimated for populations (see Edelaar & Bolnick, 2019). This way of measuring fitness can be helpful when factors that influence fitness variation change at the population level, such as environmental temperature.

Global surface temperature has increased about 0.2 °C per decade in the last 3 decades (Hansen et al., 2006) and estimates are that over the next century, temperatures will rise from 1.8 to 4.0 °C (Beaumont et al., 2011). In this scenario, it is essential to understand how temperature influences amphibian ecology. The level of vulnerability of certain species to climate change will directly depend on its thermal sensitivity (Huey et al., 2012; von May et al., 2019). There are two points to consider when thinking about the impacts of global warming: increase in mean temperatures *per se* (Thompson et al., 2013) and the increase in the magnitude of daily temperature fluctuations (IPCC, 2013; Vázquez et al., 2015). Both phenomena pose a deep concern on the future of amphibians, once they are already among the most endangered vertebrate taxa (Catenazzi, 2015; IUCN, 2018). Amphibians also have some life-history characteristics that make them even more sensitive to the consequences of climate change (Lawler et al., 2010; Katzenberg et al., 2018), such as permeable skin, limited dispersal ability, and dependence of aquatic habitats.

Here, we experimentally tested the influence of temperature on jumping performance of a Neotropical frog species, *Lysapsus limellum* Cope 1862. As different responses may have different thermal sensitivities (Kellerman et al., 2019), we measure three estimates of jumping performance. We hypothesize that more extreme temperatures will lead to a longer response time of individuals, lower first jump distance, and longer total distance travelled.

## 2. Material and Methods

### 2.1 Animals

Frogs of the genus *Lysapsus* are fully aquatic hylids that usually forage and reproduce on the surface of large water bodies in open areas (Garda et al., 2007). *Lysapsus* has a wide distribution, occurring in much of South America, which makes this genus a great study model, due to the wide variation of temperature throughout its geographic range (Sánchez et al., 2015; Cavalcanti, 2016).

Water temperatures reported at the ponds inhabited by *Lysapsus limellum* range from 26 to 32 °C (26-27.2 °C in the morning, 28.6-32 °C in the afternoon and 26.4-28 °C in the night; Garda et al., 2007). Common predators of these frogs include wolf spiders, water snakes, and giant water bugs (Garda et al., 2007; Landgref Filho et al., 2019). When threatened by predators, adults usually emit release calls and jump away, normally jumping three or four times consecutively and then diving into the water. This behavior was observed during daytime, when frogs are active (Garda et al., 2007).

### 2.2. Specimen sampling

We collected 60 adults of *Lysapsus limellum* in September of 2018 using visual encounter survey at night around ponds in the Pantanal Field Station (19° 34’ S, 57° 00’ W; DATUM=SIRGAS2000) of the Federal University of Mato Grosso do Sul, in Corumbá, Mato Grosso do Sul, central Brazil. All individuals were immediately placed into plastic bags containing water and macrophytes and transported to the laboratory, less than 1 km away.

### 2.3 Experimental design

We set up an arena with porcelain tiles measuring 170 cm in length, 70 cm in height, and 55 cm in depth, and inside it we created a thermal gradient with a 150 W infrared light at one end and ice cubes at the other, obtaining a temperature gradient ranging from 20 °C to 40 °C. In order to control the temperature inside the arena, we used thermocouples positioned at both ends and in the center of the area. Since moisture interacts with temperature influencing jumping performance of frogs (Mitchell and Bergmann, 2016), we keep the interior of the arena with a 1-cm layer of water constantly.

Each frog was used only once. We unsystematically positioned each frog in the arena, that is, the temperature to which the individual was subjected during the experiment was not predefined. Then, we used a restriction grid to maintain the frog to that temperature for ten minutes. After the acclimation, we removed the restriction grid and made a mechanical stimulus using a 30 cm cylindrical object and always by the same person at 1 m of distance. Each frog had three chances to respond to the stimulus, and between each stimulus there was an interval of 30 s. The stimulus was applied only if the individual did not respond to the previous one, resulting in a 90 s trial for each individual.

For each individual, we recorded three response variables: (1) response time, (2) distance of the first jump and, (3) total jumping distance. We recorded response time using a stopwatch, measuring the time since the first stimulus. We considered that an individual responded when it jumped, regardless of the distance. When an individual remained still for 90 s without jumping, we recorded it as an absence of response. Total jumping distance was measured with a tape as follows: at each jump, we marked the starting location and the place where the individual stopped. Then, after 90 s, we measured a straight line that the individual traveled. In all cases, temperature measurements were made only at the beginning of experiment and at the end of the 90 s period, with a digital thermometer (6802II, precision 0.1 °C).

We also measured snout-vent length (SVL) and leg length (LL) of each individual with a digital caliper (mm), and calculated the relative leg length (LL/SVL) of each individual. At the end of the experiment and morphological measurements, we safely returned all individuals to where we collected them.

### 2.4 Data analysis

To test the thermal sensitivity of each variable, we performed non-parametric regression analyses using generalized additive models (GAMs). GAMs describes the shape of complex non-linear relationships (Wood, 2001). We fitted four GAMs (using a cubic regression spline smoother) for modelling: (1) response time by a smooth of the initial temperature at which the frog was released in the arena (initial temperature, hereafter), (2) distance of the first jump by a smooth of the initial temperature, (3) total jumping distance by a smooth of the initial temperature, and (4) total jumping distance by a smooth of the final temperature reached in the arena. To account for the effect of leg length on jumping performance, we fitted one additional model with a smooth of the relative leg length for each of the four previous models. We selected the best of the two models using a Likelihood Ratio Test. If there were no difference between models, we chose the simplest one.

To test for behavioral thermoregulation, we fitted a simple linear regression relating the initial temperature with the temperature difference (final temperature minus initial temperature). All analyses were conducted in R v. 3.5.6 (R Core Team, 2019) using package mgcv (Wood, 2001).

## 3. Results

All three estimates of jumping performance were affected by temperature. The model including relative leg length and the initial temperature as main effects explained the variation in response time better than the model with only initial temperature (p = 0.004), explaining 23.6 % of the deviance. However, only the initial temperature (F_3.767_= 2.87; *P* = 0.029; Fig. 1a), and not relative leg length (F_2.000_ = 0.57, *P* = 0.572; Fig. 1b), influenced response time. The model showed that individuals under extreme temperatures (particularly at low temperatures) did not respond or increased their response time and individuals placed under mild temperatures exhibited lower response times (Fig. 1a).

**Fig. 1.**
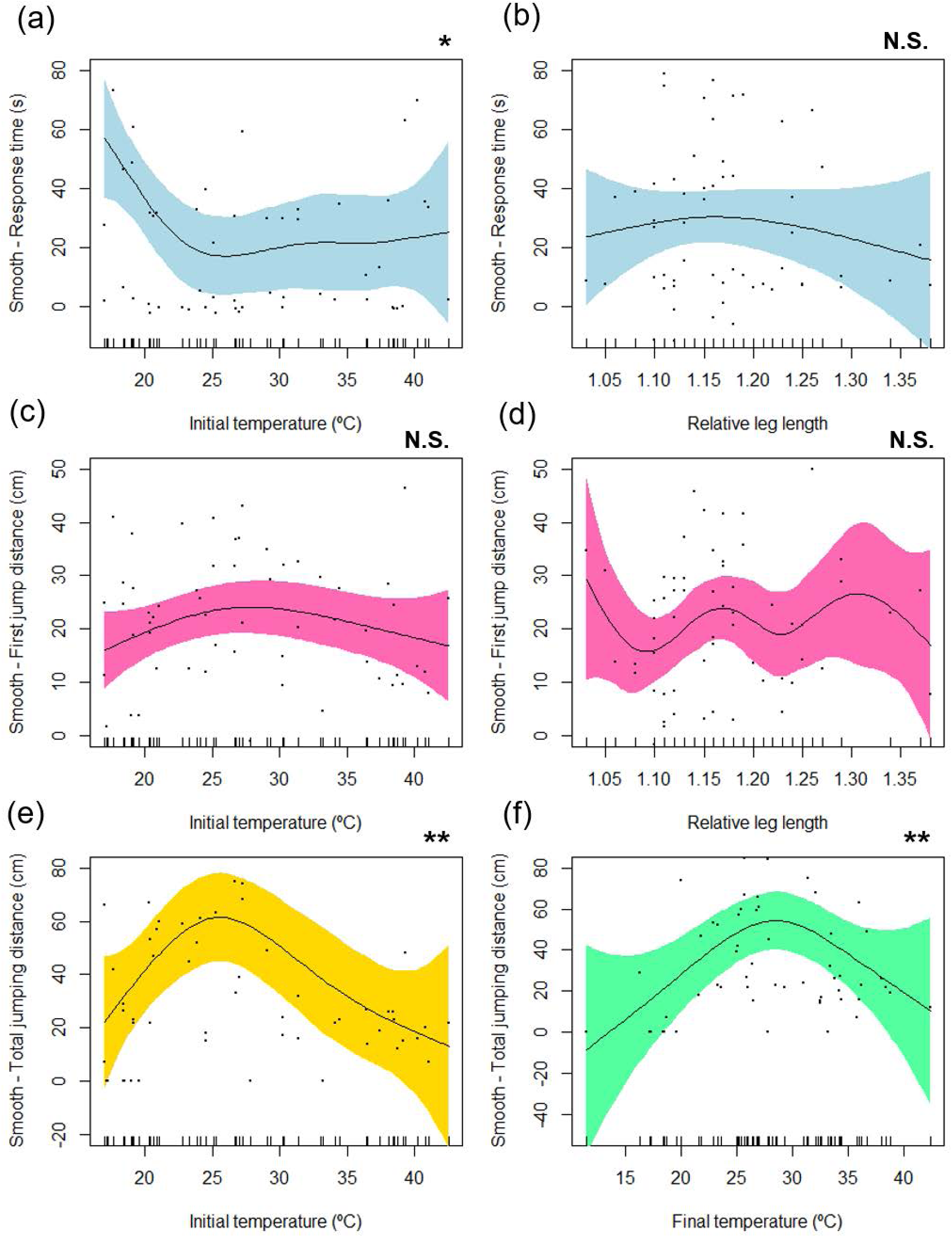
Generalized Additive Model (GAM) fits of temperature in the jumping performance of the harlequin frog, *Lysapsus limellum*. Fits of smooth functions show the additive effects (with 95% confidence intervals) of the studied predictors (** = *P* < 0.01, * = *P* < 0.05, N.S. = non-significant) on the three estimates of jumping performance: response time (s), distance of the first jump (cm), and total distance jumping distance (cm). The first model, in blue, predicted the response time of frogs by smooths of the initial temperature (a) and the relative leg length (b). The second model, in pink, predicted the distance of the first jump by smooths of the initial temperature (c) and the relative leg length (d). The third model, in yellow, revealed a significant non-linear relationship (e) between a smooth of the initial temperature and the total jumping distance. The fourth model (f) predicted the total jumping distance by a smooth of the final temperature, showing a significant non-linear relationship.

**Fig 2.**
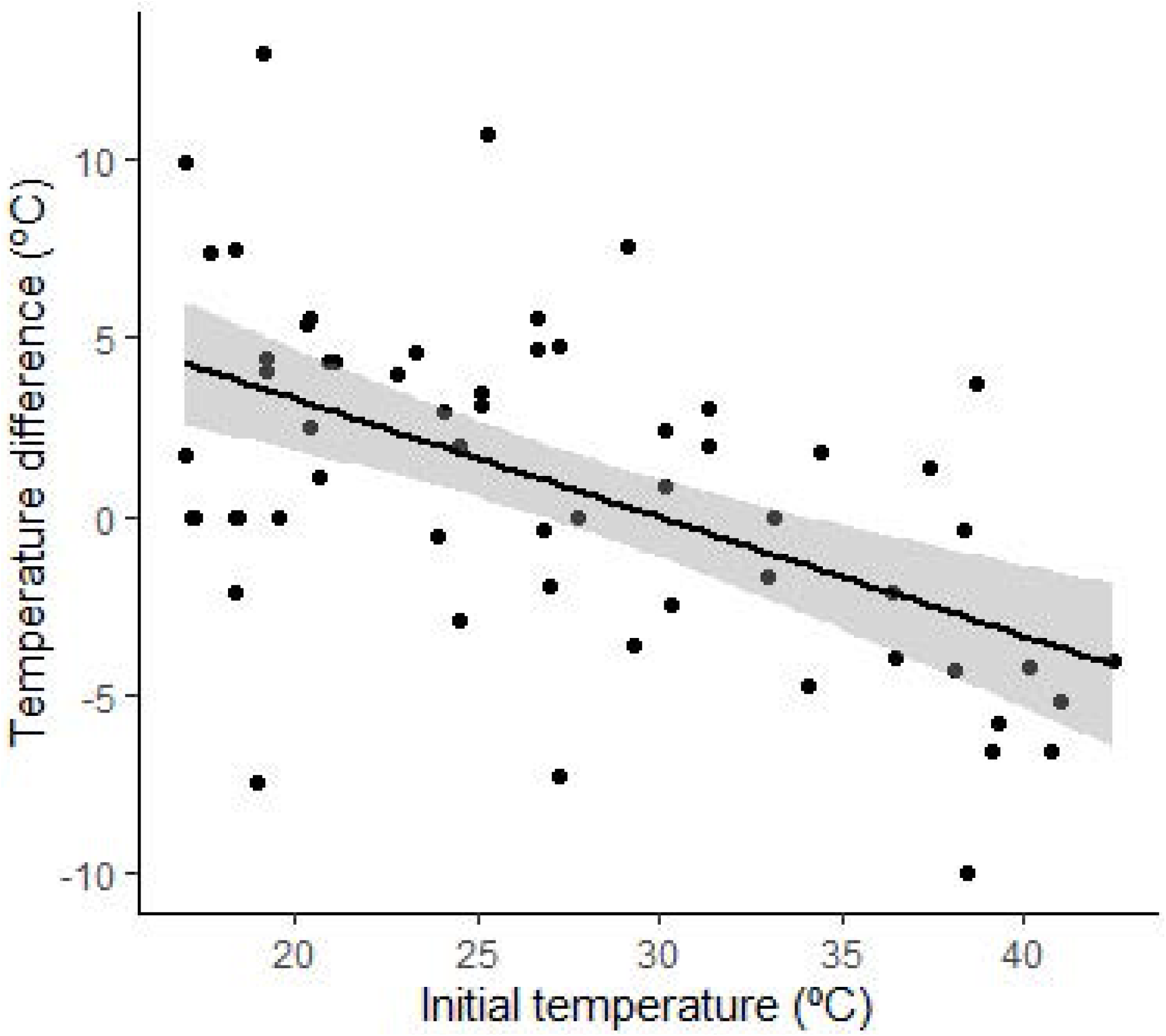
Relationship between initial temperature and temperature difference. Individuals allocated at extreme temperatures showed a larger temperature difference (F_1,58_ = 24.25; r^2^ = 0.29; *P* < 0.001).

For distance of first jump, the model including the initial temperature and relative leg length was also better than the model with only initial temperature (*P* = 0.05). This model showed that neither the initial temperature (F_2.394_ = 1.81; *P* = 0.223) (Fig. 1c), nor relative leg length (F_5.860_ = 0.98; *P* = 0.451; Fig. 1d) influenced the first jump distance. The model explained 22.1 % of the deviance.

There were no differences between models relating initial temperature and total distance jumped that included only the initial temperature and that with initial temperature and relative leg length (*P* = 0.838). The simplest model explained 19.2 % of the deviance. Total distance was influenced by the final temperature (F_2.789_ = 3.26; *P* = 0.022), being greater under extreme temperatures (Fig. 1e).

As to the relationship between the final temperature and total distance jumped, the model including relative leg length was similar to the one that did not include it (*P* > 0.05). The best model showed that final temperature significantly influenced total distance jumped (F_3.195_ = 2.95; r^2^ = 0.14; *P* = 0.038), with frogs travelling longer distances to reach milder temperatures (Fig. 1.f). The model explained 17.3 % of the deviance.

## 4. Discussion

Temperature strongly influenced the response time of frogs and the total distance jumped, while the distance of the first jump was rather constant regardless of the initial temperature. It took longer for frogs to start jumping under extreme temperatures (particularly at low temperatures), but individuals placed under extreme temperatures travelled longer distances than those in mild temperatures. Our results show that while extreme temperatures hinder jumping performance, frogs travelled long distances to find suitable thermal conditions.

We found a significant non-linear relationship between response time of jumping and the initial temperature where frogs were placed. Below approximately 23 °C frogs notably increase their response time. Response time is lowest at approximately 25 °C, which would be the optimal temperature for performance, although performance of this variable does not vary much for higher temperatures. This effect may be explained by the relationship between thermoreception and neural integration by the central nervous system, which modulates individual behavior (in our case, a latency or absence of jumping response; Abram et al., 2017). As most functions, the neural integration needed to start jumping may be related to temperature following a thermal performance curve (Little & Seebacher, 2016; Abram et al., 2017). Thus, the performance is maximum when the individual is at an environmental temperature within its optimal thermal range. As temperature becomes more extreme beyond the optimal thermal range, performance begins to be impaired, until it can no longer be performed (Huey & Stevenson, 1979). Thermal performance curves are often built at the individual level, that is, the same individual is subjected several times to different temperatures and then a performance curve is created (Careau et al., 2014). However, by measuring performance of several individuals, one can estimate the fitness component at the level of populations (Angillett,a 2006; Edelaar & Bolnick, 2019). This approach is more useful to provide estimates of the effects of climate change since they vary at the level of populations instead of individuals.

Distance of the first jump was not affected by temperature. One possible explanation for this pattern is the fleeing tactic used by this species. When threatened, individuals of this species tend to invest their energy in a greater number of short jumps instead of one or two long jumps (Garda et al., 2007). Apparently, the temperature does not seem to affect this pattern of behavior, since the distance of the first jump was not significantly different under different temperatures. Individuals of the *Pseudacris crucifer* species (also belonging to the Hylidae family) showed similar patterns in a study by Rand (1952). In this study, the author compared the jumping ability of different species, and species *Pseudacris crucifer* showed a pattern of several short jumps.

The relationship between temperature and total distance jumped was the strongest, with frogs jumping longer distances to reach places at suitable temperatures. When individuals are submitted to extreme temperatures, they tend to move to warmer places, thus increasing the difference in ratio between final and initial temperature (Mitchell & Bergman, 2016). Jumping distance is maximized at around 25 °C. At temperatures under 25 °C, jumping distance is simply reduced because locomotor response is hindered. Accordingly, our results showed that total jumping distance is maximized at final temperatures of approximately 29 °C. This temperature seems the most selected for *Lysapsus limellum*, as individuals travelled the longest distances to reach it.

As ectotherms, frogs depend on temperature to maintain their metabolic rate. These rates are maximum when animals are within their thermal tolerance limits (Halsey et al., 2015), but once these limits are exceeded, there is a risk of impairment of physiological functions (Huey & Stevenson, 1979). Travelling longer distances when submitted to thermal extremes is a trade-off (Somero, 1995; Huey & Slatkin, 1976; Vickers et al., 2011). In this trade-off, costs can be relatively high under certain circumstances. Remaining under extreme temperatures may result in muscular spasms (Paulson & Hutchison, 1987), loss of motor function (Miller & Packard, 1977), decreased fitness (Kingsolver et al., 2013), changes in growth rates (Lillywhite et al., 1973), and even death (Snyder & Weathers, 1975). Conversely, moving to a place with mild temperatures would also be costly. Firstly, because moving requires energy that could be allocated to other purposes, such as somatic growth, feeding, or reproduction (Angilletta, 2009). Searching for a better place also implies higher risk of predation (Downes, 2001). Therefore, the decision about staying in a thermally unsuitable place or moving to search for a better one probably balances costs and benefits, in order to maximize survival.

Most Harlequin frogs chose to jump in order to find mild places to park. This suggests that benefits of thermoregulating would surpass the energetic and non-energetic costs of not doing it. Mitchell and Bergmann (2016) found that *Lithobates clamitans* select places to minimize water loss, even these places being not optimal for jumping performance, and conclude that frogs hydroregulate stronger than they thermoregulate.

Taken together, our results can help understand the effects of temperature on frog performance and fitness, which will help predicting the effect of climate change on them. Neotropical frogs usually have a limited capacity for metabolic acclimation (Navas, 1996). Specifically, climate change projections are dramatic in the Pantanal. The forecast for the end of the century is that temperatures will increase around 4 °C to 7 °C and rainfall will decrease in summer and winter, promoting drier periods and greater evaporation, which will probably affect the water balance of the region (Marengo et al., 2015). All these changes can directly affect frog populations, decreasing its jumping performances, and its ability to escape predators.

## 5. Conclusions

This study demonstrates that temperature modulates the locomotor performance of *Lysapsus limellum*, following non-linear relationships, in which performance is maximized under medium temperatures and limited under extreme temperatures. Thermal sensitivity was different for two variables representing jumping performance, being stronger for the total distance moved. Although jumping was suboptimal at extreme temperatures, individuals travelled longer distances to find thermally suitable areas. This demonstrates thermoregulation capacity in these harlequin frogs.

## Acknowledgements

CCG is supported by a master’s fellowship from CNPq (GM/CNPq #133486/2018-4). ZO is a postdoctoral fellow of CAPES-PNPD (PNPD/CAPES #1694744). This study was conducted during the field course Ecology of the Pantanal, sponsored by the Ecology and Conservation Graduate Program of the Federal University of Mato Grosso do Sul (Brazil). We thank Amanda Dudczak, Jorge Dias, Camyle Maruyama for help during field experiment. Author contribution statement: CCG and DBP conceived the experiment. CCG conducted the experiment. ZO and DBP conducted data analysis and data curation, along with CCG. ZO helped interpreting the results. CCG prepared the first draft of the manuscript, to which all authors contributed further.

